# Structural insights into acetylated histone ligand recognition by the BDP1 bromodomain of *Plasmodium falciparum*

**DOI:** 10.1101/2022.08.02.501158

**Authors:** Ajit Kumar Singh, Margaret Phillips, Saleh Alkrimi, Marco Tonelli, Samuel P. Boyson, Kiera L. Malone, Jay C. Nix, Karen C. Glass

## Abstract

*Plasmodium falciparum* requires a two-host system, moving between Anopheles mosquito and humans, to complete its life cycle. To overcome such dynamic growth conditions its histones undergo various post-translational modifications to up-regulate and down-regulate certain genes required for invasion and replication. The *P. falciparum* genome encodes six bromodomain-containing proteins, of which Bromodomain Protein 1 (PfBDP1) has been shown to interact with acetylated lysine modifications on histone H3 to regulate the expression of invasion-related genes. Here, we investigated the ability of the PfBDP1 bromodomain to interact with acetyllsyine modifications on additional core and variant histones. A crystal structure of the PfBDP1 bromodomain (PfBDP1-BRD) reveals it contains the conserved bromodomain fold, but our comparative analysis between the PfBDP1-BRD and the 8 human bromodomain families indicates it has a unique binding mechanism. Solution NMR spectroscopy and ITC binding assays carried out with acetylated histone ligands demonstrate that it preferentially recognizes tetra-acetylated histone H4, and we detected weaker interactions with multi-acetylated H2A.Z in addition to the previously reported interactions with acetylated histone H3. Our findings indicate PfBDP1 may play additional roles in the *P. falciparum* life cycle, and the distinctive features of its bromodomain binding pocket could be leveraged for the development of new therapeutic agents to help overcome the continuously evolving resistance of *P. falciparum* against currently available drugs.

## 1. INTRODUCTION

Eukaryotic cells accommodate nearly three meters of genetic information into a relatively small space within the nucleus by wrapping ~146 bp of DNA around an octamer of histone to form the nucleosome [1–3]. In addition to this highly ordered packaging, the high affinity interactions between the histone proteins and DNA restricts the accessibility of the genetic information by various accessory proteins required to perform essential cell functions such as DNA replication, recombination, repair, and transcription [4–6]. The histone octamer is made up of the H3-H4 histone tetramer and two H2A-H2B histone dimers [2, 3, 7]. The protein sequence of histone H3 and H4 are highly conserved among all species ranging from yeast to humans, whereas the protein sequence of H2A and H2B histones can vary greatly [8]. The histone proteins are highly basic in charge, and contain a globular C-terminal region along with a protease sensitive N-terminal tail that protrudes from the surface of nucleosome core [9]. The flexible N-terminal tail region of the histone protein is abundant in lysine, arginine, and serine residues, which undergo various covalent post-translational modifications (PTMs) such as acetylation, phosphorylation, methylation, ubiquitination, and SUMOylation [6, 9]. The histone tail PTMs are known to modulate the DNA-nucleosome interaction, and are also recognized by various accessory proteins required for gene activation and/or gene silencing [10]. Thus, the combinations of PTMs are also referred to as a histone code, which is an important epigenetic mechanism responsible for regulating access to the genetic information [5, 9, 11, 12]. The enzymes responsible for managing the type and position of these specific PTMs, are known as writers (methylases, acetylases, kinases, and ubiquitinases), and erasers (demethylases, deacetylases, phosphatases) of the code [9, 13]. There are also protein reader modules (bromodomains, chromodomains, WD40 repeat, and PHD fingers, etc.) that recognize the epigenetic signals provided by particular PTMs and recruit associated proteins/cellular machinery to genomic positions to carry out their function [14]. For example, bromodomains, which are known to recognize acetylated lysine residues on the N-terminal histone tails, share an evolutionarily conserved structural fold comprised of four alpha-helices connected by two variable-length loops, which form the acetyllysine binding pocket [15]. In humans, there are approximately 60 human bromodomains divided into eight subfamilies, and they are often found in chromatin remodeling proteins and transcription factors that are involved in essential cellular functions including transcription, DNA replication, and DNA repair [15].

Bromodomain-containing proteins have also been identified in parasitic organisms including *Plasmodium falciparum* [16]. This unicellular eukaryote exhibits a very complex life cycle for its survival and replication. It requires a two-host system, moving between the Anopheles mosquito (invertebrate) and humans (vertebrate), and invades multiple cell types such as insect cells, hepatocytes, and red blood cells [17]. To adapt to such dynamic environmental conditions its genome undergoes various post-translational modifications to regulate the specific activation or inactivation of genes at particular stages [18]. The genome of the *Plasmodium falciparum* is organized by the four canonical core histones (H3, H4, H2A, and H2B), as well as variant histones H2A.Z, H2B.V, H3.3, and CenH3 [19–21]. These histones are reported to be enriched with acetylated lysine residues in the transcriptionally active euchromatin region. Using mass spectrometry techniques, several acetylated lysine residues have been identified on the N-terminal tail region of *P. falciparum* histone proteins including H3 (K4ac, K9ac, K14ac, K18ac), H4 (K5ac, K8ac, K12ac, K16ac, K20ac), H2A.Z (K11ac, K15ac, K19ac), and H2B.Z (K3ac, K8ac, K13ac, K14ac, K18ac) [19, 20, 22, 23]. Some of these modifications are known to occur at specific stages of its life cycle, namely histone H3K9ac/H3K14ac, and their function has been linked to the activation of invasion-related genes required for entry into blood cells [24]. The *P. falciparum* genome encodes histone acetyltransferases (HATs), histone deacetylases (HDACs), and histone methyltransferases (HMTs) that dynamically regulate the presence of specific PTMs [25, 26]. The *P. falciparum* genome also encodes for ten bromodomain-containing proteins namely, Bromodomain Proteins 1-7 (PfBDP1, PfBDP2, PfBDP3, PfBDP4, PfBDP5, PfBDP6, PfBDP7, PfBDP8, GCN5 (histone acetyltransferase GCN5), and SET1 (SET domain protein 1)), which likely recognize acetylated lysine modifications. Recent studies on PfBDP1 have shown that it is a multi-domain protein with a predicted ankyrin repeat in the N-terminal region, and a bromodomain (BRD) near the C-terminus [16, 24, 27–29]. PfBDP1 is thought to play an important role in the activation of invasion-related genes in the asexual stage as its knockdown drastically affects the replication of *P. falciparum* in red blood cells. The bromodomain of PfBDP1 was shown to recognize specific acetylated lysine marks, such as histone H3K9ac and H3K14ac, over methylated or unmodified histone H3 [24]. In addition, histone peptide pull-down assays also indicate that PfBDP1 can interact with acetyllysine modifications on histone H4, and the histone variants H2B.Z and H2B.A [22]. Interestingly, PfBDP1 appears to form a core complex with two other bromodomain-containing proteins, PfBDP2 and PfBDP7, which were also enriched with acetylated histones in the pull-down assays [22, 24, 27]. PfBDP7 is an essential gene in *P. falciparum* that was recently shown to co-localize with the genome-wide binding sites of PfBDP1. Both proteins are commonly found at the promoters of invasion genes, however, PfBDP1 and PfBDP7 are also enriched in heterochromatin at genes encoding the variant surface antigens, which contribute to parasite infection. Thus, they are also thought to function as an important regulatory complex for silencing the expression of variant surface antigen [27]. Similarly to the SAGA and RSC chromatin remodeling complexes, which contain multiple bromodomain proteins, PfBDP7 has been shown to interact with PfBDP1 and PfBDP2, likely allowing the complex to recognize a specific subset of acetylation marks on histone proteins [22, 24, 27].

Among the five *Plasmodium* species, *Plasmodium falciparum* is the largest contributor to malaria infections in humans. According to the 2019 WHO report, nearly half of the world’s population is at risk of malaria, which includes around 229 million active cases and 409,000 deaths annually. The most affected groups from malaria includes children below the age of five. While the development of a vaccine to prevent this deadly disease shows great promise, human mortality is still predicted to increase [30, 31]. This is due in part to low vaccination rates in young children, and also due to evolutionary processes that drive parasite resistance against the currently available antimalarial drugs [32]. Thus, new therapeutics will continually be needed to treat infected patients. A better understanding of the *Plasmodium falciparum* life cycle, and the molecular mechanisms driving infection and disease progression are needed to develop additional anti-malarial strategies. Bromodomains are validated drug targets for the treatment of various diseases [33]. Additional structural and functional studies on the PfBDP1 bromodomain will provide valuable information in search of small molecule inhibitors against it.

We hypothesized that the PfBDP1-BRD would recognize multiple acetylation modifications, similarly to many of the human BRD counterparts. As such, we examined the structural fold of the PfBDP1-BRD and carried out a systematic investigation of the bromodomain binding activity with acetylated histone ligands. We used X-ray crystallography to structurally characterize the PfBDP1-BRD, and *in vitro* Isothermal Titration Calorimetry (ITC) and Nuclear Magnetic Resonance (NMR) binding studies, to investigate its ability to recognize specific acetyllysine modifications on histones H3, H4, H2A.Z, and H2B.Z. Our results outline the structural differences between the PfBDP1 bromodomain compared to human bromodomains, and we characterized its functional activity by identifying its preferred histone ligands. Our results also provide new insights into the potential role(s) of PfBDP1 in the life cycle of *Plasmodium falciparum*. Together, our structurefunction approach will aid in the design of specific therapeutics to inhibit the activity of PfBDP1, which may offer a mechanism to restrict the replication of *P. falciparum* in red blood cells to treat the disease.

## 2. RESULTS

### 2.1. Crystal structure of the PfBDP1-BRD

Bromodomains are conserved reader domains that are known to recognize acetyllysine marks on the N-terminal tails of histone proteins. Prior research on the PfBDP1-BRD indicates that it plays an important role in recognizing acetylated histone H3 to regulate the expression of invasion-related genes, which facilitates parasite entry into red blood cells. The PfBDP1 protein is organized into multiple domains consisting of seven ankyrin repeats at the N-terminus and a bromodomain towards the C-terminal region (residues 333-456) (**Fig. 1A**). However, there is no information available on the molecular mechanisms utilized by the PfBDP1-BRD to coordinate histone ligand recognition and binding. To structurally characterize the PfBDP1-BRD in complex with the histone H3K14ac ligand, an expression construct containing residues 333-456 was used for the crystallization trials. Protein crystals of PfBDP1-BRD (333-456) grown in a complex with the H3K14ac ligand diffracted to 2.0 Å. The structure was solved by molecular replacement with a monomer of the PfBDP1 bromodomain in the asymmetric unit. The structure was refined to a final Rwork and Rfree values of 19.49% and 23.65%, respectively, and the data collection and refinement statistics are shown in **Table 1**. Disappointingly, no electron density was observed for the histone H3K14ac ligand. However, the final structure of PfBDP1-BRD (333-456) protein possesses the conserved bromodomain fold consisting of four left-handed alpha helices (**Fig. 1B).** The four-helix bundle is arranged in two halves connected by the AB and BC loops (residues 389-393 and 413-417). The first half is comprised of the aZ (333-357 aa) and aA (379-389 aa) helices. The end of the aZ helix encodes a ‘HIF’ shelf that forms the bottom of the binding pocket, and the long and variable ZA loop (358-378 aa) frames one side of the binding pocket before connecting to the aA helix. Part of the ZA loop also forms a short a-helix from residues 369-374, which is a conserved structural feature of the bromodomain binding pocket [15]. The second half of the conserved bromodomain fold is comprised of the aB helix (394-413 aa) and the aC helix (417-455 aa) that form the other side of the binding pocket. The aB helix is 19 aa shorter than the aC helix and is connected by a 3-residue long BC loop, which contains the conserved asparagine residue N413 that is involved in the coordination of the acetyllysine moiety (358-378 aa) (**Fig. 1B**). Thus, the PfBDP1 bromodomain possesses the characteristic structural features that enable the recognition and binding of acetyllysine post-translational histone modifications.

**Fig. 1.**
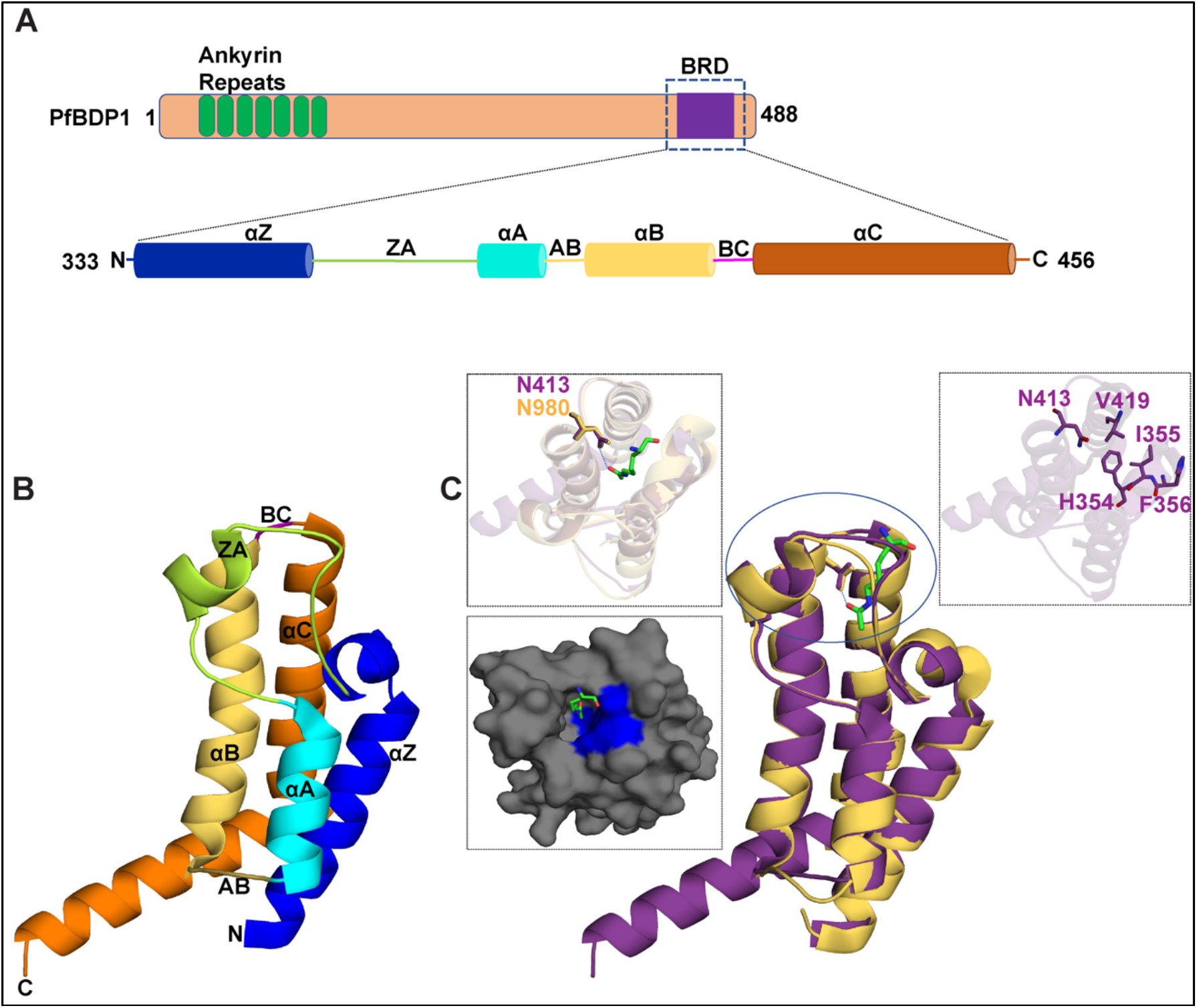
Structural features of the PfBDP1 protein. (A) Domain organization of PfBDP1 protein, where the ankyrin repeats are highlighted in green, loop regions in tan, and the bromodomain (BRD) in purple. The BRD is further expanded to show the amino acid sequence and highlights the basic amino acids in blue, acidic amino acids in red, neutral amino acids in black, and hydrophobic amino acids in gray. The secondary structure assigned to regions of the amino acid sequence is based on the 2.0 Å crystal structure of the PfBDP1-BRD (333-456) (PDB ID: 7M97). Hydrophobic amino acids found in the binding pocket are marked with an asterisk, while the acidic and basic residues in the binding pocket are annotated by a diamond. (B) Cartoon representation of the PfBDP1-BRD (333-456) (PDB ID: 7M97) structure, where the aZ helix is blue, the aA helix is cyan, aB helix is yellow, aa helix is orange, the ZA loop is green, and the BC loop is magenta. (C) Structural alignment of PfBDP1-BRD (333-456) in purple with the TIF1α bromodomain (PDBID: 3O35) in yellow. The acetyllysine found in the TIF1α bromodomain is colored green and represented in stick model. The left panel insert shows a zoomed in view of the binding pockets and coordination of acetyllysine by the TIF1α bromodomain and the surface representation of the predicted binding pocket of PfBDP1-BRD (333-456) where the hydrophobic residues are highlighted in blue and the acetyllysine group from the overlaid TIF1 a bromodomain structure is in green. The right panel insert shows the conserved N413 and gatekeeper residues V419 and HIF motif (354-356 aa) in stick model.

**Table 1.** Data collection and structure refinement statistics for the PfBDP1-BRD.

|  |  |
| --- | --- |
| **Data collection (PDB ID: 7M97)** |  |
| No. of crystals | 1 |
| Beamline | ALS 4.2.2 |
| Wavelength | 0.9762 |
| Detector | RDI CMOS 8M |
| Crystal to detector distance (mm) | 225 |
| Rotation range per image | 0.2 |
| Total rotation range (°) | 180 |
| Exposure time per image (s) | 0.5 |
| Resolution range (Å) | 34.97 – 2.0 (2.072-2.000) |
| Space group | I 2 2 2 |
| **Unit cell parameters** |  |
| a,b,c (Å) | 38.051, 88.667, 106.797 |
| $\alpha,\beta,\gamma$ (°) | 90, 90, 90 |
| Unique reflections | 12597 (1203) |
| R<sub>work</sub>/R<sub>free</sub> (%) | 19.49/23.65 |
| Multiplicity | 7.0 (6.4) |
| Mean I/s (I) | 19.24 (2.93) |
| Completeness (%) | 99.63 (99.25) |
| R<sub>merge</sub> (%) | 2.602 (26.72) |
| Wilson B factor (Å) | 29.36 |
| **Number of non-hydrogen atoms** |  |
| Macromolecules | 1006 |
| Solvent | 110 |
| Protein residues | 124 |
| **RMSD** |  |
| Bond length (Å) | 0.008 |
| Bond angles (°) | 089 |
| **Ramachandran plot** |  |
| Favored regions (%) | 98.36 |
| Allowed regions (%) | 1.64 |
| Outliers (%) | 0.00 |

### 2.2 Comparison of the structural features of the PfBRD1 binding pocket with the human TIF1α-BRD/ TRIM24 PHD-BRD

Bromodomains have been identified to specifically recognize the acetylation marks by their conserved asparagine and gatekeeper residues. To determine these residues in PfBDP1-BRD its structure was compared with the human bromodomains in a complex with an acetylated histone ligand. PfBDP1-BRD shows the closest structural similarity with TIF1α-BRD (TRIM24 PHD bromodomain) acetyllysine complex (PDB ID: 3O35) with an RMSD (root mean square deviation) value of 0.88. This this structure was used for depicting the PfBDP1-BRD acetyllysine binding pocket. Like other bromodomains, the predicted acetyllysine binding pocket of PfBDP1-BRD (333-456) has a conserved asparagine residue (Asn413) responsible for making a hydrogen bond to the acetyllysine, while the gatekeeper residue valine (Val419) and the HIF shelf motif (res 354-356) are important for ligand binding specificity **(Fig. 1C)**. Similar to human bromodomains, it also has an abundance of hydrophobic residues, where six hydrophobic residues were present on the aZ, aB, aC helices, and 4 hydrophobic residues were present on the ZA loop **(Fig. 1C)**. In conclusion, our structural analysis indicates PfBDP1-BRD can also specifically recognize acetyllysine marks on histone proteins.

### 2.3 Comparison of PfBDP1-BRD with human bromodomains

To further define the unique features of the PfBDP1-BRD, we carried out an in-depth structural analysis to compare the similarities and differences of the PfBDP1-BRD with the structures of human bromodomains. In humans, there are 61 bromodomains that are classified into eight families based on a structural alignment. We compared our structure of the PfBDP1-BRD to one member from each family that was selected based on the highest sequence identity and the availability of a deposited crystal structure in PDB. Thus, we used the structure of the CECR2 BRD from family I, the second BRD from BRD4 in family II, the CREBBP BRD in family III, the ATAD2B BRD in family IV, the TIF1α BRD from family V, the TRIM28 BRD in family VI, the second BRD from TAF1 in family VII, and the third BRD from PB1 in family VIII (3). Interestingly, the PfBDP1-BRD has the highest sequence identity (48.44%) with the family III BRD of CREBBP, while it had the lowest sequence identity with the family VI BRD of TRIM28 (22.95%) (**Table 2 and Suppl. Fig. 2A**).

**Table 2.**
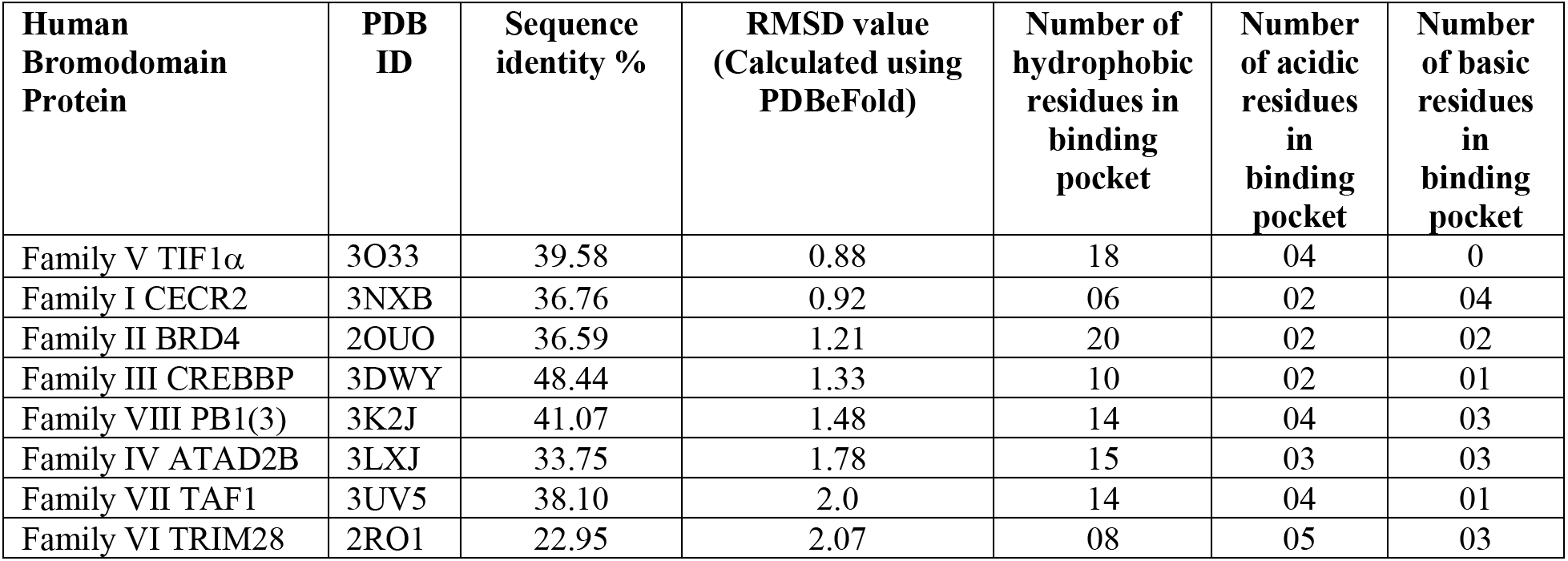
Structural analysis of the PfBDP1-BRD (PDB ID 7M97) compared to human bromodomains in families I-VIII.

| **Human Bromodomain Protein** | **PDB ID** | **Sequence identity %** | **RMSD value (Calculated using PDBeFold)** | **Number of hydrophobic residues in binding pocket** | **Number of acidic residues in binding pocket** | **Number of basic residues in binding pocket** |
| --- | --- | --- | --- | --- | --- | --- |
| Family V TIF1α | 3O33 | 39.58 | 0.88 | 18 | 04 | 0 |
| Family I CECR2 | 3NXB | 36.76 | 0.92 | 06 | 02 | 04 |
| Family II BRD4 | 2OUO | 36.59 | 1.21 | 20 | 02 | 02 |
| Family III CREBBP | 3DWY | 48.44 | 1.33 | 10 | 02 | 01 |
| Family VIII PB1(3) | 3K2J | 41.07 | 1.48 | 14 | 04 | 03 |
| Family IV ATAD2B | 3LXJ | 33.75 | 1.78 | 15 | 03 | 03 |
| Family VII TAF1 | 3UV5 | 38.10 | 2.0 | 14 | 04 | 01 |
| Family VI TRIM28 | 2RO1 | 22.95 | 2.07 | 08 | 05 | 03 |

To investigate the structural similarities and differences of PfBDP1-BRD with each of these human bromodomain proteins, we first calculated the RMSD using the PDBeFold online server with our PfBDP1-BRD crystal structure as the query structure [34]. The structural fold of PfBDP1-BRD shows the highest similarity with the BRD of CECR2 in family I, having an RMSD value of only 0.88. In contrast, the largest structural differences were observed between the PfBDP1-BRD and the family IV BRD of ATAD2B, which displayed an RMSD value of 2.13 (**Table 2 and Suppl. Fig. 2B**).

The crystal structure of PfBDP1-BRD was also compared with the human bromodomains to understand the differences in amino acid composition in the acetyllysine binding pocket with a particular focus on hydrophobic and charged amino acid residues, which are known to play an important role in the recognition of acetyllysine residues. As shown in **Fig. 2A and Table 2** the PfBDP1-BRD (PDB ID: 7M97) has 10 hydrophobic residues in the acetyllysine binding pocket, which is similar to the number of hydrophobic residues found in the CREBBP BRD. Whereas it has fewer hydrophobic residues than bromodomains of BRD4, KIAA1240, TIF1α, TAF1, and PB1(3), and more than the CECR2 and TRIM28 human bromodomains. The electrostatic surface potential map (**Fig. 2B**) shows the PfBDP1-BRD is most similar to the ATAD2B and CECR2 BRDs with 6 charged amino acid residues (acidic and basic) in the acetyllysine binding pocket. Its binding pocket appears to be more highly charged than the BRDs of BRD4, CREBBP, TIF1α, and TAF1, and less charged in comparison to the TRIM28 and PB1(3) BRDs (**Table 2 and Fig. 2B**). Our systematic analysis of the BRD binding pockets illustrates that besides having a close structural similarity with human bromodomains, PfBDP1-BRD has a unique chemical composition in the acetyllysine binding pocket, which may help in designing a specific drug inhibitor against PfBDP1-BRD.

**Fig. 2.**
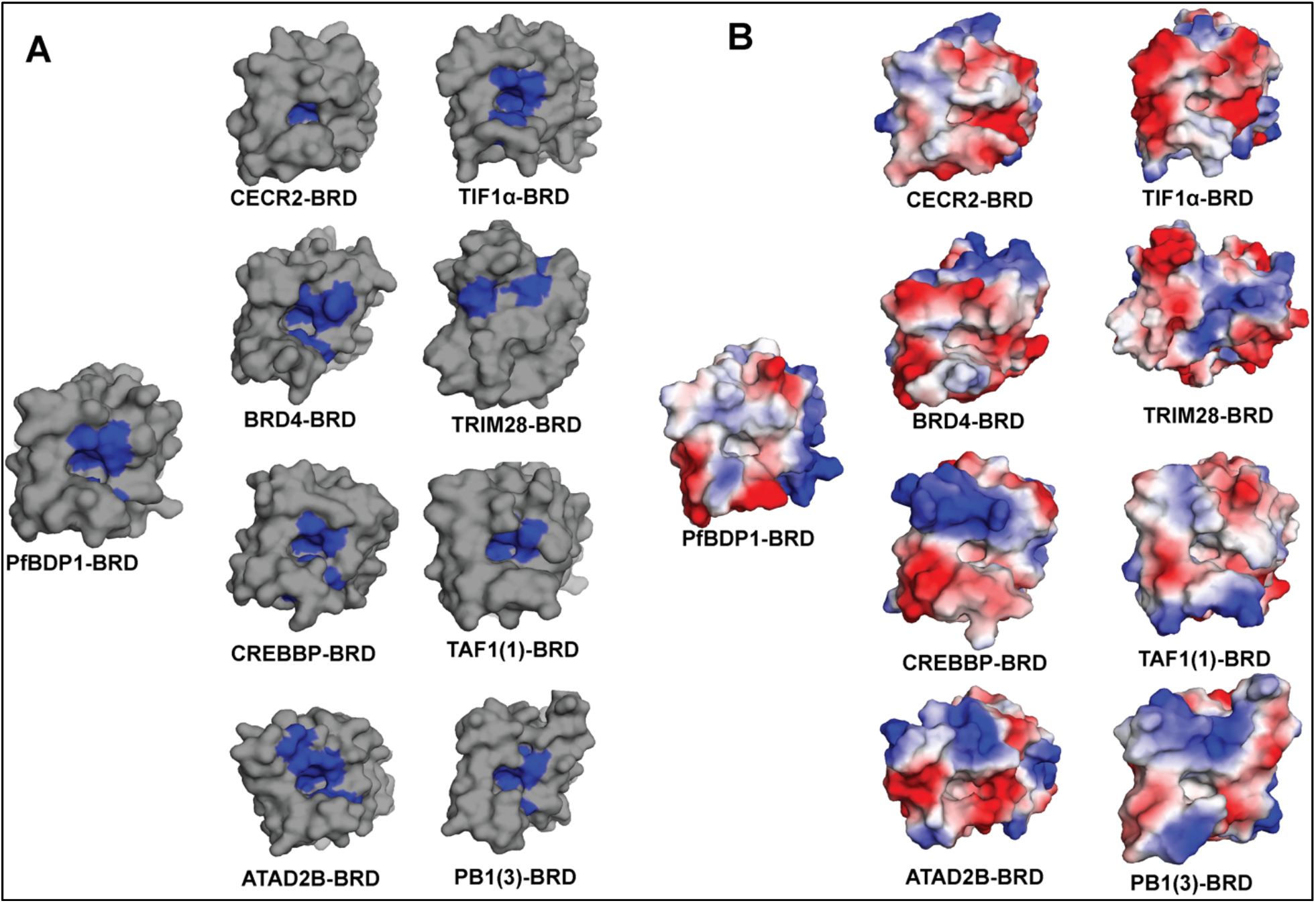
Comparison of hydrophobic and electrostatic features of the BRD binding pockets. (A) Surface representation of the PfBDP1-BRD (331-456) is depicted alongside representative human bromodomains from families I-VIII. The blue regions in the left panel highlight the hydrophobic residues near the acetyllysine binding pocket, while the gray region shows the overall surface of the bromodomains. (B) An electrostatic surface potential map was generated using PyMOL for the PfBDP1-BRD (331-456) and representative human bromodomains. Electrostatic surface potentials are shown over a range of −88.229kT/e (red) and +88.229kT/e (blue).

### 2.4 The PfBDP1-BRD preferentially interacts with acetylated histone H4 ligands

Bromodomains are known to play a major role in decoding the epigenetic information by recognizing acetyllysine modifications to recruit bromodomain-containing proteins and any associated proteins/complexes to the chromatin [35]. The PfBDP1-BRD was previously shown to interact with histone H3 acetylated at lysine 9 and 14 (H3K9ac and H3K14ac), and this activity was required for the activation of invasion related genes in *Plasmodium falciparum* [24]. However, characterization of the specific binding interactions of the PfBDP1-BRD with PTM histone ligands was only carried out with a limited number of unmodified, acetylated, and methylated lysine residues on histone H3 [24]. Human bromodomains are frequently able to recognize multiple acetylation modifications on the core histones. For example, the well-studied BET bromodomain family binds to single and multiple acetyllysine mark combinations on histones H3 and H4 [15, 35, 36]. Based on the observed structural similarities between the PfBDP1-BRD and human bromodomains as described above, we hypothesized that it would likely interact with additional acetyllysine modifications on both histone H3 and H4. To investigate this, we carried out isothermal titration calorimetry (ITC) binding assays with a library of *P. falciparum* histone peptides that contain acetyllysine modifications at all possible positions within the N-terminal tail of each histone. As shown in **Table 3** we confirmed the interaction of PfBDP1 BRD with the diacetylated H3K9acK14ac (K_D_ = 1125 ± 181 μM), and mono-acetylated H3K14ac (K_D_ = 1215.0 ± 162.3 μM) histone ligands by ITC. However, the isothermal entropy plots showed no binding to the H3K9ac or the unmodified histone H3 ligands (**Table 3, Fig. 3B, Suppl. Fig. 3**). On the other hand, when the binding activity of PfBDP1-BRD was tested against histone H4 peptides monoacetylated at lysine residue K5, K8, K12, K16, and K20, it was able to recognize the H4K5ac (K_D_ 1210 ± 208 uM) and H4K12ac (K_D_ 1070 ± 158 uM) ligands, but did not interact with H4K8ac, H4K16ac, H4K20ac, or the unmodified H4 ligands. Interestingly, the PfBDP1-BRD is able to select for di-acetylated H4 ligands including H4K5acK8ac (K_D_ = 1510.0 ± 168.0 μM) and H4K5acK12ac (K_D_ = 621.0 ± 28.3 μM), but it demonstrated the strongest binding affinity with the tetra-acetylated histone H4 peptide (H4K5acK8acK12acK16ac, K_D_ = 117.0 ± 11.3 μM) (**Table 3, Fig. 3B, Suppl. Fig. 3**).

**Fig. 3.**
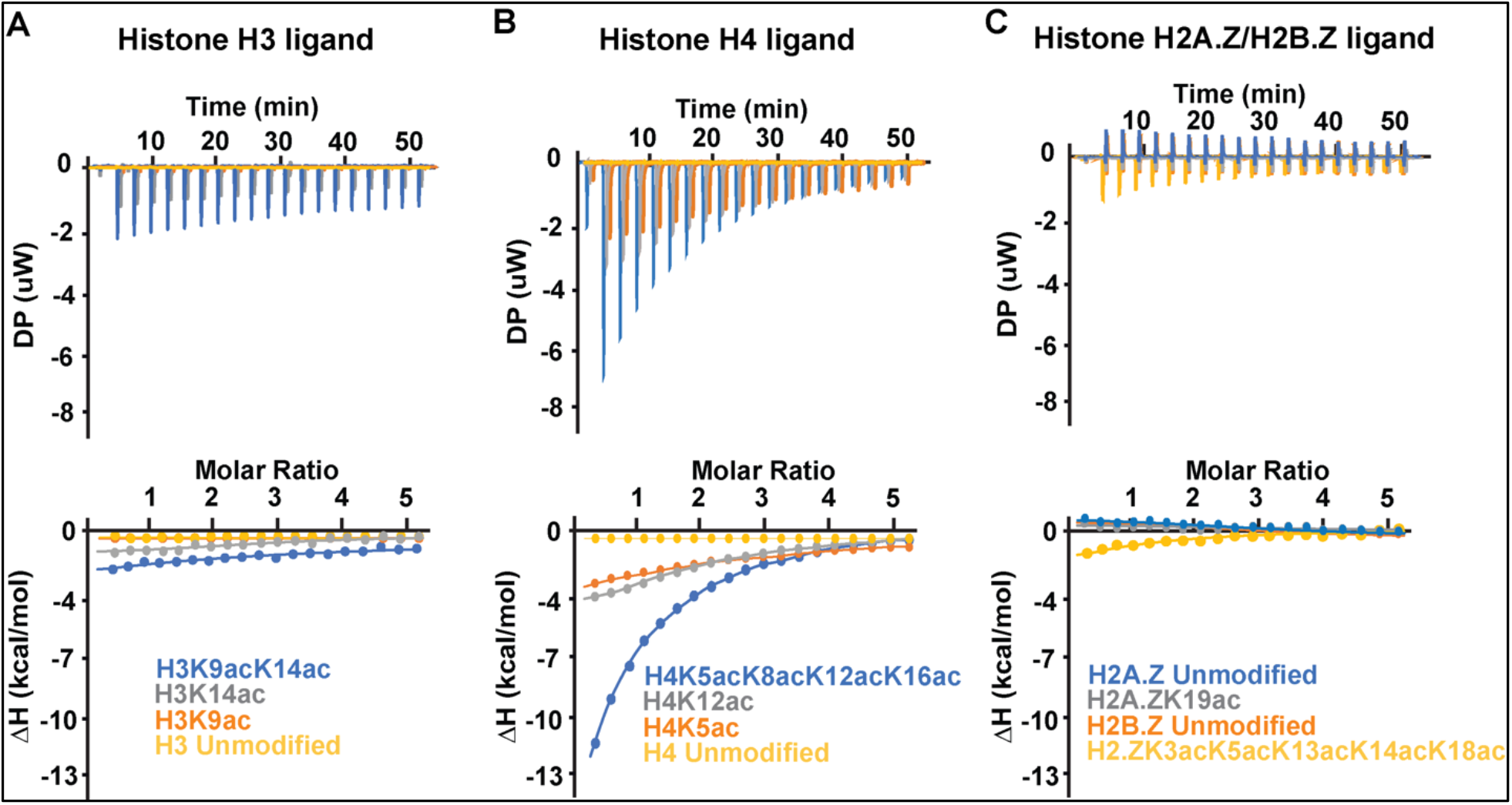
Superimposed ITC enthalpy plots for the binding of PfBDP1-BRD with acetylated histone ligands. (A) PfBDP1-BRD with histone H3 peptide ligands, where unmodified H3 is shown in yellow, H3K9ac in orange, H3K14ac in gray and H3K9acK14ac in blue. (B) PfBDP1-BRD with histone H4 peptide ligands, where the unmodified H4 is shown in yellow, H4K5ac in orange, H4K12ac in gray, and H4K5acK8acK12acK16ac in blue. (C) PfBDP1-BRD with the histone H2A.Z and H2B.Z peptide ligands, where unmodified H2A.Z is shown in blue, H2A.ZK19ac in gray, H2B.Z unmodified in orange, and H2B.ZK3acK8acK13acK14acK18ac in yellow.

**Table 3.**
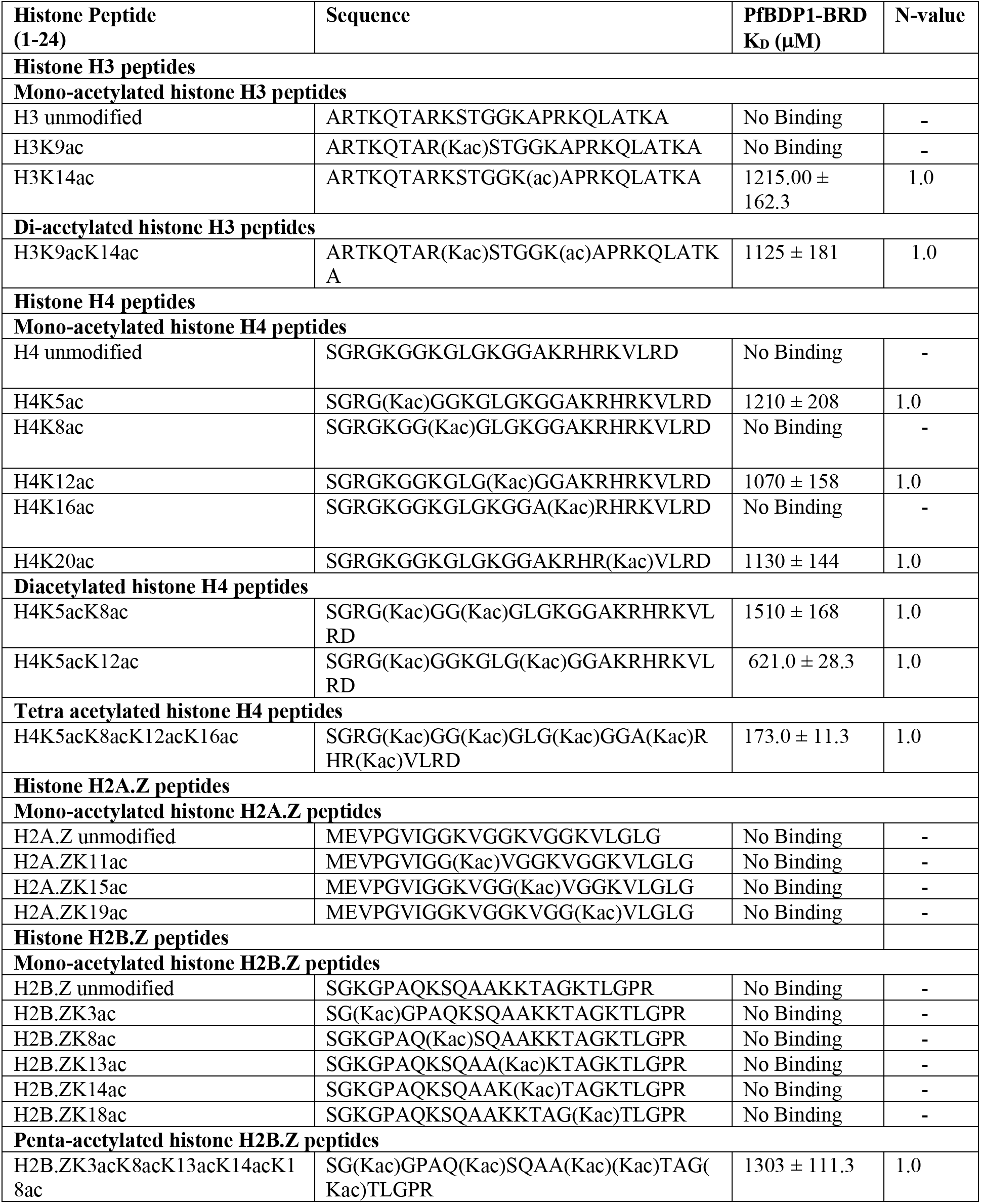
Binding affinities of the PfBDP1-BRD with acetylated histone peptides measured by ITC.

A recent study by Hoeijmakers, et. al., used histone peptide pull-downs coupled to mass spectrometry to identify histone readers and associated proteins in nuclear extracts from blood-stage *P. falciparum* parasite cultures [22]. They discovered that the PfBDP1-BRD was enriched at acetyllysine modifications on the parasite-specific tails of histone variants H2A.Z and H2B.Z [22]. Thus, we carried out additional ITC binding assays to further evaluate the interaction of PfBDP1-BRD with histone H2A.Z containing acetylated lysine at residues K11, K15, and K19, and with histone H2B.Z acetylated at positions K3, K8, K13, K14, and K18. However, in our study, we did not observe any interactions of the PfBDP1-BRD with the mono-acetylated histone variants H2A.Z and H2B.Z. To further examine if multiple acetyllysine modifications may be required to recruit the PfBDP1-BRD to histone variants we also tested its ability to bind the penta-acetylated H2B.ZK3acK8acK13acK14acK18ac, and it does show a weak interaction with this ligand (**Table 3, Fig. 3B, Suppl. Fig. 3**). Taken together, our results indicate that the PfBDP1-BRD recognizes specific acetylated lysine marks found on histones H3 and H4, and preferentially binds to histone H4 when it contains multiple acetyllysine residues. This suggests that effector-mediated recognition of the histone tails by PfBDP1-BRD is dependent on the combinatorial readout of multiple post-translational modifications over a span of amino acids on the histone tails.

To further confirm the binding of acetylated histone peptides to the PfBDP1-BRD we used NMR to carry out chemical shift perturbation studies (CSPs). We recorded two-dimensional ^15^N-^1^H heteronuclear single quantum coherence (HSQC) spectra and examined the changes in chemical shift of apo-bromodomain backbone amide peaks upon addition of unmodified and acetylated histone H4, H3, H2A.Z, and H2B.Z peptides. The absence of CSPs upon the addition of unmodified histone peptides to the PfBDP1-BRD indicates no interaction (**Suppl. Fig. 4**). whereas distinct CSPs observed for the PfBDP1-BRD residues in the presence of acetylated histone peptides confirm the binding interactions observed by ITC (**Fig. 4**). Among the acetylated histone H4 peptides (residues 1-24) tested, the tetra acetylated H4K5acK8acK12acK16ac showed the most significant CSPs. This is supported by our ITC data, which shows binding of K_D_ = 173.0 ± 11.3 μM, and is higher than the binding affinities for H4K5ac (K_D_ = 1210 ± 208 μM) and H4K12ac (K_D_ = 1070 ± 158 μM). Similar to the unmodified histone H4, there were no CSPs observed upon the addition of the unmodified histone H3 peptide (**Suppl. Fig. 4**). Interestingly, although the binding affinity of H3K9acK14ac was stronger than the H3K14ac ligand, we did not observe any CSPs in the presence of the singly modified histone H3K9ac ligand. This is complementary to our ITC data which also demonstrates no binding for the H3K9ac peptide. The addition of H3K14ac histone peptide-induced only weak CSPs. We also confirmed no binding to the H2A.ZK11ac ligand as illustrated by the absence of CSPs. However, we did observe weak binding of the H2B.ZK3acK8acK13acK14acK18ac histone ligand as illustrated by the presence of CSPs in the NMR spectra. In conclusion, our NMR titration experiments supplement our ITC data and confirm that the histone H4K5acK8acK12acK16ac peptide binds with the highest affinity to PfBDP1-BRD, followed by the H4K12ac, H3K14ac, and the H2B.ZK3acK8acK13acK14acK18ac histone peptides.

**Fig. 4.**
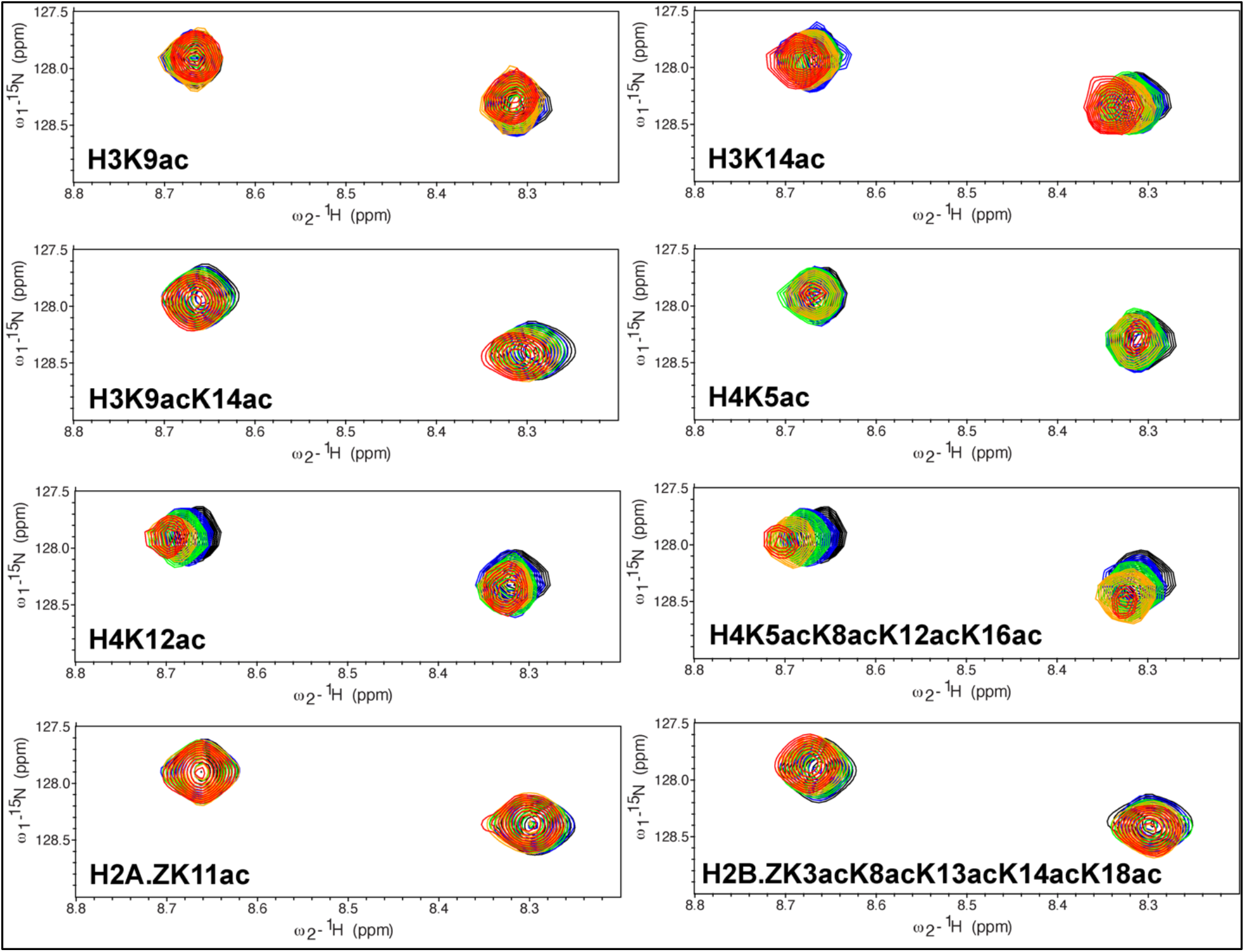
Interaction of the PfBDP1-BRD with acetylated histone ligands. Superimposed 2D ^15^N-^1^H HSQC spectra of the ^15^N-labeled PfBDP1-BRD collected in titration experiments with the indicated histone peptides. The peaks of the PfBDP1-BRD apo protein are shown in black. The titration of PfBDP1-BRD with histone peptides was performed at increasing concentrations of each ligand and include 1:0.5 molar ratio (blue), 1:1 molar ratio (green), 1:2.5 molar ratio (orange) and in 1:5 molar ratio (red). Each spectrum is labeled with the histone peptide used for the titration.

## 3. DISCUSSION

In this study, we determined the highest resolution crystal structure deposited for the PfBDP1-BRD, and confirm it contains the conserved bromodomain fold and a functional acetyllysine binding pocket. Importantly, at this resolution, we have also observed continuous density for the ZA loop, which makes up an important part of the acetyllysine binding pocket that was missing in the previously deposited structure (**Suppl. Fig. 1**). Our ITC and NMR binding studies of PfBDP1-BRD with acetylated histone ligands indicate it can specifically recognize the acetyllysine modifications on histones H3, H4, and H2B.Z. We demonstrated that the PfBDP1-BRD binds to mono- and di-acetylated histone H3K14ac and H3K9acK14ac, respectively. These results are similar to what was previously reported by Josling et. al. [24], however, we did not observe any interaction with the mono-acetylated histone H3K9ac ligand. Furthermore, our binding studies identified additional interactions of the PfBDP1-BRD with histone H4 ligands including, monoacetylated (H4K5ac, H4K12ac, H4K20ac), di-acetylated (H4K5acK8ac, H4K5acK12ac), and tetra-acetylated (H4K5acK8acK12acK16ac). Mass spectrometry experiments conducted by Hoeijmakers et. al. suggested that PfBDP1 interacts with acetylation marks on histone H2B.Z [22]. Our binding studies indicate that the PfBDP1-BRD does not interact with monoacetylated H2A.Z and H2B.Z ligands, but the ITC and NMR titration assay support a weak interaction that may exist with the penta-acetylated histone H2B.ZK3acK8acK13acK14acK18ac ligand. To date PfBDP4-BRD in complex with an acetylated histone peptide (PDBID 5VS7) is the only ligand bound crystal structure available for *P. falciparum* bromodomains. From our interaction studies, we identified the PfBDP1-BRD recognizes the tetra-acetylated histone H4 peptide with the highest affinity. Hence, further investigations need to be conducted to understand how these additional acetylation marks are recognized by PfBDP1 on nucleosomes. Since humans are one of the primary hosts for *Plasmodium falciparum*, which in turn causes malaria, we carried out a detailed analysis comparing the PfBDP1-BRD with 61 bromodomains identified in the humans [15]. PfBDP1-BRD shows high structural identity with the human bromodomains, indicating it likely performs a similar function to its human counterparts. We found that although the PfBDP1-BRD can be structurally aligned with the human bromodomains with an RMSD of less than 2.1, it has a low sequence identity (>50%) with these proteins. However, the significant differences in primary sequence between PfBDP1-BRD and human bromodomains, suggest that the PfBDP1-BRD utilizes a unique molecular mechanism to select for specific acetyllysine modifications on the *P. falciparum* histones. For example, PfBDP1-BRD has the highest structural identity with the TIF1α-BRD (RMSD = 0.88), but only 39.58% sequence identify. Ligand binding studies on the TIF1α-BRD show that it binds to the acetylated histone peptide H4K16ac with a K_D_ of 92.6 ± 1.9 μM by ITC, whereas the PfBDP1-BRD does not show any interaction with the H4K16ac mark [15]. In addition, the PfBDP1-BRD has low structural (RMSD = 2.0) and sequence identity (38.10%) with the TAF1-BRD(1). As expected the TAF1-BRD(1) does not have any overlapping binding activity with the PfBDP1-BRD [15]. Differences in the primary sequences between the PfBDP1-BRD and human bromodomains contribute to the unique features of the acetyllysine binding pocket. Our structural analysis revealed that the number of hydrophobic residues and surface electrostatics of the PfBDP1-BRD has some similarities with the CREBBP and TRIM28 human bromodomains, but it seems to contain an exclusive combination of ligand specificity determinants. For example, the conserved acetyllysine anchor in the PfBDP1-BRD is close to the gatekeeper residue at the top of the aC helix. The small hydrophobic Val gatekeeper residue in PfBDP1-BRD is conserved with the family II BET-BRDs, which also have the ability to select for multiply acetylated histones [15]. However, instead of the WPF shelf residues found in the family II BED-BRDs, PfBDP1-BRD has a HIF shelf that is only observed in the human family IV BRPF3-BRD. Thus, the PfBDP1-BRD appears to be a cross between two human bromodomain families by having a ‘wide’ acetyllysine binding pocket observed in the BET bromodomains, combined with the shelf motif from BRPF3, which has a ‘keyhole’ type pocket due to its large Phe gatekeeper residue [15, 37]. This distinctive arrangement of specificity determinants in the binding pocket, in combination with a unique set of hydrophobic and electrostatic features will be important drivers underlying the molecular mechanism of ligand recognition by the PfBDP1-BRD. It also provides a valuable opportunity to design small molecule inhibitors that can selectively target the PfBDP1-BRD via the rational design of chemical-based interactions optimized to take advantage of these features.

Bromodomain-containing proteins are known to play a major role in the regulation of gene activation or inactivation and have been validated as a potent therapeutic target for the treatment of various diseases such as cancer, neurological disorders, and inflammation [33]. Currently, there are several small-molecule bromodomain inhibitors in clinical trials against cancer, and they may be useful for treating neurological disorders due to their ability to cross the highly selective bloodbrain barrier [38]. Several small-molecule human bromodomain inhibitors have also been used in targeting *P. falciparum* bromodomains. For example, the triazolophthalazine-based small molecule L-45 shows very strong affinity for the PfGCN5-BRD, and its co-crystal structure highlights residues in the acetyllysine binding pocket that are conserved between the parasite bromodomain and its human ortholog. Another small molecule BI-2536, which is an approved PLK1 and JAK2-FLT3 kinase inhibitor, binds with high affinity to the PfBDP3-BRD [28, 29]. Interestingly, a small molecule inhibitor (JIB-04 E) was discovered to be active towards the *P. falciparum* Jumonji histone lysine demethylase (PfJmj3). JIB-04 E was able to mimic the transcriptional changes observed upon knockdown of PfBDP1 by downregulating several invasion-related gene families [39]. To date there are not any effective small molecule inhibitors available that target the PfBDP1-BRD, although a weak binding interaction with SGC-CBP30 was observed in docking studies [40]. Additionally, a recent preprint reports *in silico* binding studies suggesting that the compound, MCULE-6567089130, could be a potential PfBDP1-BRD inhibitor [41]. Thus, the new data obtained in this study enables a further understanding of the structure and function of PfBDP-BRD and will be a valuable resource in the search for small molecule inhibitors using a similar approach employed to target human bromodomains.

The PfBDP1 protein plays an essential role in the *P. falciparum* life cycle and survival, supporting the idea that this protein could be an effective anti-malarial drug target [16, 24, 29]. With resistance quickly developing against antimalarial drugs, it has become increasingly important to understand the etiology and molecular mechanisms driving malaria infections in order to identify additional drug targets for the development of new antimalarials. An approach using a combination of anti-malarial drugs that target multiple life cycle stages of *P. falciparum* would likely increase the efficacy of treatment while also reducing the development of resistance. The identification of target proteins that have a significant role in various stages of life cycle of *P. falciparum* may provide additional therapeutic utility. Histone PTMs are known to change dynamically during different stages of the cell cycle, throughout development, and as disease progresses [42], the interaction of the PfBDP1-BRD may be important for regulating additional stages of the *P. falciparum* life-cycle. Our study establishes an additional activity of the PfBDP1-BRD in recognizing histone H4 and H2B.Z acetylation marks, whose functional significance with respect to the regulation of gene expression in *P. falciparum* has yet to be explored. The PfBDP1 protein has also been shown to form a multi-subunit complex with PfBDP2 and/or PfBDP7 [24, 27, 43]. In addition to the C-terminal bromodomain, the full-length PfBDP1 protein contains 7 ankyrin repeats located near the N-terminus, which may function as a protein-protein interaction domain. Thus, the interaction of the bromodomain with acetylated H3, H4, and H2B.Z histones would recruit PfBDP1 to chromatin, and the ankyrin repeats could serve as a bridge between PfBDP1 and PfBDP2/PfBDP7, which would allow them to work together in a complex to regulate gene activation and inactivation processes. Our studies suggest that PfBDP1 could perform important chromatin regulatory functions at various stages of the *P. falciparum* life cycle and provide additional utility as a target in the development of new approaches for anti-malarial therapeutics.

Although nine bromodomain-containing proteins have been identified in *P. falciparum*, functional studies are available on only three of these proteins including, PfBDP1, PfGCN5, and PfBDP7 [24, 27, 44]. The domain organization illustrates that all of them except PfBDP3 have a C-terminal bromodomain, which is located on the N-terminus, and that PfBDP4 has two consecutive bromodomains in the C-terminal region [45]. A phylogenetic analysis of bromodomains in Protozoan parasites indicates that PfBDP1, PfBDP2, and PfBDP3 cluster into an evolutionarily related family, which suggests that PfBDP2/3 could perform functions similar to PfBDP1 [16]. Furthermore, PfBDP1 is evolutionary and most closely related to the *Toxoplasma gondii* Bromodomain Protein 1 (TgBDP1) [16]. As a functional ortholog, TgBDP1 may have similar biological roles in the development of disease. Thus, our current study outlining the structure and function of PfBDP1-BRD may also translate to the TgBDP1-BRD and assist in the design of bromodomain inhibitors that specifically target both *T. gondii* and *P. falciparum*.

## 4. MATERIALS AND METHODS

### 4.1 Plasmid construction

The DNA sequence encoding PfBDP1-BRD (333-456) region was amplified using polymerase chain reaction (PCR) taking PfBDP1 full length as a template. The PCR amplified PfBDP1-BRD (333-456) region was subsequently cloned into the pDEST15 vector containing the N-terminal glutathione S-transferase (GST) tag using the Gateway cloning approach. The codon-optimized PfBDP1-BRD (303-488) construct cloned in a pGEX-6P-1 vector with an N terminal GST-tag followed by a precision protease site was obtained from GenScript.

### 4.2 Protein overexpression and chromatographic purification

The PfBDP1 BRD (333-456) and PfBDP1 BRD (303-488) constructs in a pGEX-6P-1 vector were transformed into *E. coli* BL21 (DE3) pLysS cells and allowed to grow in a TB medium containing 100 μg/ml ampicillin and 25 μg/ml chloramphenicol at 37 °C at OD_600_ 0.8. The cells were induced with 0.2 mM isopropyl β-D-thiogalactopyranoside (IPTG) and induction was allowed to proceed for 16 hrs at 20°C. The cells were harvested by centrifugation at 5,000 RPM for 10 mins. The cell pellet was homogenized in a lysis buffer containing 50 mM HEPES (pH 7.5), 500 mM NaCl, 1 mM EDTA, 2 mM DTT, 10%Glycerol, 0.1mg/ml lysozyme, and a protease inhibitor tablet (Thermo Scientific). The homogenized cells were lysed by sonication and centrifuged at a speed of 11,000 RPM for 60 mins at 4°C to pellet the insoluble material. The obtained supernatant was incubated at 4°C for 2h with 10 mL of Glutathione Beads packed in a 25 mL chromatography column (G Biosciences). After incubation beads were washed with 200 mL wash buffer containing 20 mM HEPES (pH 7.5), 500 mM NaCl, 1 mM EDTA, and 2 mM DTT followed by GST-tag cleavage using Precision protease at 4°C for 14h. After GST cleavage, the PfBD1 BRD proteins were eluted with wash buffer. SDS-PAGE gels stained with GelCode Blue (Thermo Scientific) were used to confirm the purity of the PfBDP1 BRD protein samples. The protein concentration was determined by absorbance at 280 nm using a NanoDrop (Thermo Scientific) and an extinction coefficient of 18910 M^-1^cm^-1^ for PfBDP1 BRD (333-456) and 29910 M^-1^cm^-1^ for PfBDP1 BRD (303-488). For X-ray crystallization experiments the protein was further purified by size-exclusion chromatography (SEC) using a HiPrep 16/60 Sephacryl S-100 HR column at 4 °C in a buffer containing 20 mM HEPES (pH 7.5), 150 mM NaCl, and 2 mM DTT.

### 4.3 Histone peptide synthesis

All the unmodified and acetylated histone peptides were purchased from GenScript (Piscataway, NJ, USA). The synthesized peptides were purified by HPLC to achieve 98% purity and present in lyophilized form with TFA salt. The identity of each peptide was confirmed with mass spectrometry and the sequence of the peptides with modifications are listed in **Table 3.** Despite multiple attempts the *P. falciparum* H2A.ZK11acK15acK19ac could not be synthesized.

### 4.4 Isothermal Titration calorimetry

Isothermal titration calorimetry (ITC) experiments were carried out using a MicroCal PEAQ-ITC and a MicroCal iTC200 (Malvern Panalytical). The purified PfBDP1 BRD proteins (containing residues 303-488 or 333-456) were dialyzed for 48 hrs. at 4°C into ITC buffer containing 20 mM sodium phosphate buffer pH 7.5, 150 mM NaCl, and 1 mM TCEP. The dialyzed PfBDP1 BRD (303-488 or 333-456) proteins were added to the sample cell at concentrations of 100-200 uM, and between 2.5 – 5.0 mM of each histone peptide in the ITC dialysis buffer was placed in the injection syringe. A total of 20 injections were carried out, with the parameters set as follows. The first injection had a volume of 0.5 uL, a reaction time of 0.4 sec, followed by a 180-sec delay. The remaining 19 injections were completed with an injection volume of 2 uL, a reaction time of 4 sec, and a 150 sec spacing time between each injection. All ITC runs were carried out at 5°C with a differential power (DP) value of 10, the feedback set to high, at a stirring speed of 750 RPM, and the with an initial delay for the reaction set to 60 sec.

The ITC data obtained were analyzed using the MicroCal PEAQ-ITC 200 analysis software provided with the instrument (Malvern Panalytical). The heat of dilutions for the buffer and peptide ligands were obtained from titrating buffer into buffer, buffer into PfBDP1 BRD protein (303-488 and 333-456), and histone peptide into buffer. The heats of dilution for the histone peptides were subtracted from the raw isotherms obtained from the histone peptide and PfBDP1 BRD (303-488 and 333-456) protein titrations. The first injection was discarded from the data analysis, and the data was integrated using a 1:1 binding model assuming a single set of identical binding sites. A Marquandt nonlinear least-squares analysis was used to calculate the binding affinities and N values from the normalized heat changes. Binding experiments were repeated in triplicate for each ligand and the mean K_D_ was calculated from the average of the three runs, with the standard deviation coming from the mean. Non-binding ligand experiments were repeated in duplicate.

### 4.5 Nuclear Magnetic Resonance Spectroscopy

Chemical shift perturbation experiments were performed using 0.3 - 0.5 mM of uniformly ^15^N-labeled PfBDP1-BRD (303-488) in NMR buffer containing 20 mM Tris-HCl pH 6.8, 150 mM NaCl, 10 mM DTT and 10 % D2O. Titration mixtures of the PfBDP1-BRD (303-488) and each of the modified histone peptides (res 1-24) unmodified H3, H3K9ac, H3K14ac, H3K9acK14ac, unmodified H4, H4K5ac, H4K12ac, H4K5acK8acK12acK16ac, unmodified H2A.Z, H2A.ZK11ac, unmodified H2B.Z, H2B.ZK3acK8acK13acK14acK18ac were prepared at concentrations of 1:0, 1:0.5, 1:1, 1:2.5, and 1:5 molar ratio, in a total volume of 80 μL, The sample mixtures were then transferred into 1.7-mm NMR tubes (Bruker). Two-dimensional ^15^N-^1^H HSQC (heteronuclear single quantum coherence) experiments for all samples were acquired on Bruker Avance III spectrometer equipped with a cryogenic 1.7mm probe. The temperature of the samples was regulated at 25°C throughout data collection. All spectra were processed using TopSpin (Bruker) as described previously [46] and analyzed using NMRFAM-SPARKY [Lee W, Tonelli M, Markley JL. Bioinformatics. 2015 Apr 15; 31(8):1325-7].

### 4.6 Crystallization of PfBDP1 BRD (333-456)

For crystallization, the PfBDP1-BRD (333-456) was concentrated to ~15 mg/mL in a buffer containing 20 mM HEPES (pH 7.5), 150 mM NaCl and 2 mM DTT. The crystallization trials were done using commercially available conditions such as Index HT Screen (Hampton Research), Crystal Screen HT (Hampton Research), JSCG-plus HT-96 (Molecular Dimension), and the Ligand Friendly Screen HT-96 (Molecular Dimension). Crystals were set up in a 96-well sitting drop plate (96-well Swissci 3-well midi UVXPO crystallization plate, HR3-125 Hampton research) with a reservoir volume of 40 μL and a drop volume of 2 μL. The PfBDP1 BRD (333-456) protein at 10 mg/mL with 5.3 mM of the histone H3K14ac (res 1-19) ligand was mixed in ratios of either 1:1 or 1:3 with the mother liquor and grown using a sitting drop vapor diffusion method at 4°C.

Initial PfBDP1-BRD (333-456) protein crystals were obtained after three days of incubation at 4°C in condition D7 from the JCSG-plus HT-96 crystal screen 0.2 M Lithium sulfate, 0.1 M Tris pH8.5, 40% v/v PEG400, 100 μL 1,8-ANS. The crystal conditions were optimized in a 24-well VDX hanging drop tray (Model/company) at 4°C. The final crystals grew in a 2 μL drop containing 1 μL of 12 mg/mL PfBDP1 BRD mixed with 5.3 mM of histone H3K14ac (res 1-19) ligand, and 1 μL of mother liquor containing 0.3 M lithium sulfate, 0.1 M Tris-HCL pH 8.5, and 38% PEG400. The crystal was harvested in a 75 μm Dual Thickness Microloop LD with 18 mm pins inserted into a B1A (ALS style) reusable goniometer base from MiTeGen (Ithaca, NY, USA). The crystals were flash frozen in liquid nitrogen at 100 K. A high-resolution dataset was collected at the Advanced Light Source (ALS) beamline 4.2.2 and recorded on an RDI CMOS-8M detector. The data were processed using XDS [47], and an initial structure solution was obtained by molecular replacement using PHASER with PDBID: 5ULC as the search model [48]. Iterative rounds of model building, and refinement were carried out with PHENIX and COOT [49], and the final model of the PfBDP1 BRD was deposited into the PDB under PDBID: 7M97. The structure figures and structural superpositions were prepared using PyMOL [50].

## PDB deposition

The coordinates and structure factors of the PfBDP1 BRD (residues 333-456) structure was deposited in the Protein Data Bank (PDBID: 7M97).

## CRediT authorship contribution statement

A.K.S., M.P., S.A., M.T., S.P.B., K.L.M., and J.C.N. performed the experiments and analyzed the data together with K.C.G. A.K.S prepared all the final figures. A.K.S and K.C.G. wrote the manuscript with edits from all authors.

## Acknowledgments and funding sources

Special thanks to ACPHS PharmD student Brittany Allen who’s passion for infectious disease started us along this scientific journey. Plasmids for expression of the full-length PfBDP1 protein were kindly provided by Dr. Michael Duffy at the University of Melbourne. This study made use of the Advanced Light Source in Berkley, CA and the National Magnetic Resonance Facility at Madison (NMRFAM). Beamline 4.2.2 of the Advanced Light Source, a DOE Office of Science User Facility under Contract No. DE-AC02-05CH11231 is supported in part by the ALS-ENABLE program funded by the National Institutes of Health, National Institute of General Medical Sciences, grant P30 GM124169-01. NMRFAM is supported by NIH grant R24GM141526. Automated DNA sequencing was performed in the Vermont Integrative Genomics Resource DNA Facility and was supported by the University of Vermont Cancer Center, Lake Champlain Cancer Research Organization, and the UVM Larner College of Medicine. Research reported in this publication was supported by an Institutional Development Award (IDeA) from the National Institute of General Medical Sciences of the National Institutes of Health under grant number P20GM103449. Its contents are solely the responsibility of the authors and do not necessarily represent the official views of NIGMS or NIH. This research was also supported by an ACPHS graduate research assistantship and Blythe award to SA, and by the UVM Larner College of Medicine.

## REFERENCES

[1] K.E. Van Holde, C.G. Sahasrabuddhe, B.R. Shaw, A model for particulate structure in chromatin, Nucleic Acids Res 1(11) (1974) 1579–86.

[2] K. Luger, A.W. Mader, R.K. Richmond, D.F. Sargent, T.J. Richmond, Crystal structure of the nucleosome core particle at 2.8 A resolution, Nature 389(6648) (1997) 251–60.

[3] R.K. McGinty, S. Tan, Nucleosome structure and function, Chem Rev 115(6) (2015) 2255–73.

[4] A.A. Amodeo, D. Jukam, A.F. Straight, J.M. Skotheim, Histone titration against the genome sets the DNA-to-cytoplasm threshold for the Xenopus midblastula transition, Proc Natl Acad Sci U S A 112(10) (2015) E1086–95.

[5] B.D. Strahl, C.D. Allis, The language of covalent histone modifications, Nature 403(6765) (2000) 41–5.

[6] D.E. Sterner, S.L. Berger, Acetylation of histones and transcription-related factors, Microbiol Mol Biol Rev 64(2) (2000) 435–59.

[7] A.J. Andrews, K. Luger, Nucleosome structure(s) and stability: variations on a theme, Annu Rev Biophys 40 (2011) 99–117.

[8] S. Sullivan, D.W. Sink, K.L. Trout, I. Makalowska, P.M. Taylor, A.D. Baxevanis, D. Landsman, The Histone Database, Nucleic Acids Res 30(1) (2002) 341–2.

[9] T. Kouzarides, Chromatin modifications and their function, Cell 128(4) (2007) 693–705.

[10] A.J. Bannister, T. Kouzarides, Regulation of chromatin by histone modifications, Cell Res 21(3) (2011) 381–95.

[11] T. Jenuwein, C.D. Allis, Translating the histone code, Science 293(5532) (2001) 1074–80.

[12] S.K. Zaidi, D.W. Young, M. Montecino, J.B. Lian, J.L. Stein, A.J. van Wijnen, G.S. Stein, Architectural epigenetics: mitotic retention of mammalian transcriptional regulatory information, Mol Cell Biol 30(20) (2010) 4758–66.

[13] L.A. Baker, C.D. Allis, G.G. Wang, PHD fingers in human diseases: disorders arising from misinterpreting epigenetic marks, Mutat Res 647(1-2) (2008) 3–12.

[14] R.K. Prinjha, J. Witherington, K. Lee, Place your BETs: the therapeutic potential of bromodomains, Trends Pharmacol Sci 33(3) (2012) 146–53.

[15] P. Filippakopoulos, S. Picaud, M. Mangos, T. Keates, J.P. Lambert, D. Barsyte-Lovejoy, I. Felletar, R. Volkmer, S. Muller, T. Pawson, A.C. Gingras, C.H. Arrowsmith, S. Knapp, Histone recognition and large-scale structural analysis of the human bromodomain family, Cell 149(1) (2012) 214–31.

[16] V. Jeffers, C. Yang, S. Huang, W.J. Sullivan, Jr., Bromodomains in Protozoan Parasites: Evolution, Function, and Opportunities for Drug Development, Microbiol Mol Biol Rev 81(1) (2017).

[17] W.C. Lee, B. Russell, L. Renia, Sticking for a Cause: The Falciparum Malaria Parasites Cytoadherence Paradigm, Front Immunol 10 (2019) 1444.

[18] J. Tang, S.A. Chisholm, L.M. Yeoh, P.R. Gilson, A.T. Papenfuss, K.P. Day, M. Petter, M.F. Duffy, Histone modifications associated with gene expression and genome accessibility are dynamically enriched at Plasmodium falciparum regulatory sequences, Epigenetics Chromatin 13(1) (2020) 50.

[19] M.B. Trelle, A.M. Salcedo-Amaya, A.M. Cohen, H.G. Stunnenberg, O.N. Jensen, Global histone analysis by mass spectrometry reveals a high content of acetylated lysine residues in the malaria parasite Plasmodium falciparum, J Proteome Res 8(7) (2009) 3439–50.

[20] J. Miao, Q. Fan, L. Cui, J. Li, J. Li, L. Cui, The malaria parasite Plasmodium falciparum histones: organization, expression, and acetylation, Gene 369 (2006) 53–65.

[21] W.A. Hoeijmakers, A.M. Salcedo-Amaya, A.H. Smits, K.J. Francoijs, M. Treeck, T.W. Gilberger, H.G. Stunnenberg, R. Bartfai, H2A.Z/H2B.Z double-variant nucleosomes inhabit the AT-rich promoter regions of the Plasmodium falciparum genome, Mol Microbiol 87(5) (2013) 1061–73.

[22] W.A.M. Hoeijmakers, J. Miao, S. Schmidt, C.G. Toenhake, S. Shrestha, J. Venhuizen, R. Henderson, J. Birnbaum, S. Ghidelli-Disse, G. Drewes, L. Cui, H.G. Stunnenberg, T. Spielmann, R. Bartfai, Epigenetic reader complexes of the human malaria parasite, Plasmodium falciparum, Nucleic Acids Res 47(22) (2019) 11574–11588.

[23] A. Saraf, S. Cervantes, E.M. Bunnik, N. Ponts, M.E. Sardiu, D.W. Chung, J. Prudhomme, J.M. Varberg, Z. Wen, M.P. Washburn, L. Florens, K.G. Le Roch, Dynamic and Combinatorial Landscape of Histone Modifications during the Intraerythrocytic Developmental Cycle of the Malaria Parasite, J Proteome Res 15(8) (2016) 2787–801.

[24] G.A. Josling, M. Petter, S.C. Oehring, A.P. Gupta, O. Dietz, D.W. Wilson, T. Schubert, G. Langst, P.R. Gilson, B.S. Crabb, S. Moes, P. Jenoe, S.W. Lim, G.V. Brown, Z. Bozdech, T.S. Voss, M.F. Duffy, A Plasmodium Falciparum Bromodomain Protein Regulates Invasion Gene Expression, Cell Host Microbe 17(6) (2015) 741–51.

[25] A.M. Salcedo-Amaya, W.A. Hoeijmakers, R. Bartfai, H.G. Stunnenberg, Malaria: could its unusual epigenome be the weak spot?, Int J Biochem Cell Biol 42(6) (2010) 781–4.

[26] N. Coetzee, H. von Gruning, D. Opperman, M. van der Watt, J. Reader, L.M. Birkholtz, Epigenetic inhibitors target multiple stages of Plasmodium falciparum parasites, Sci Rep 10(1) (2020) 2355.

[27] J.E. Quinn, M.D. Jeninga, K. Limm, K. Pareek, T. Meissgeier, A. Bachmann, M.F. Duffy, M. Petter, The Putative Bromodomain Protein PfBDP7 of the Human Malaria Parasite Plasmodium Falciparum Cooperates With PfBDP1 in the Silencing of Variant Surface Antigen Expression, Front Cell Dev Biol 10 (2022) 816558.

[28] H.H.T. Nguyen, L.M. Yeoh, S.A. Chisholm, M.F. Duffy, Developments in drug design strategies for bromodomain protein inhibitors to target Plasmodium falciparum parasites, Expert Opin Drug Discov 15(4) (2020) 415–425.

[29] C. Tallant, P. Bamborough, C.W. Chung, F.J. Gamo, R. Kirkpatrick, C. Larminie, J. Martin, R. Prinjha, I. Rioja, D.F. Simola, R. Gabarro, F. Calderon, Expanding Bromodomain Targeting into Neglected Parasitic Diseases, ACS Infect Dis 7(11) (2021) 2953–2958.

[30] N. Praet, K.P. Asante, M.C. Bozonnat, E.J. Akite, P.O. Ansah, L. Baril, O. Boahen, Y.G. Mendoza, V. Haine, S. Kariuki, M. Lamy, K. Maleta, R. Mungwira, L. Ndeketa, A. Oduro, B. Ogutu, F. Olewe, M. Oneko, M. Orsini, F. Roman, E.R. Bahmanyar, D. Rosillon, L. Schuerman, V. Sing’oei, D.J. Terlouw, S. Wery, W. Otieno, J.Y. Pircon, Assessing the safety, impact and effectiveness of RTS,S/AS01E malaria vaccine following its introduction in three sub-Saharan African countries: methodological approaches and study set-up, Malar J 21(1) (2022) 132.

[31] D.J. Weiss, A. Bertozzi-Villa, S.F. Rumisha, P. Amratia, R. Arambepola, K.E. Battle, E. Cameron, E. Chestnutt, H.S. Gibson, J. Harris, S. Keddie, J.J. Millar, J. Rozier, T.L. Symons, C. Vargas-Ruiz, S.I. Hay, D.L. Smith, P.L. Alonso, A.M. Noor, S. Bhatt, P.W. Gething, Indirect effects of the COVID-19 pandemic on malaria intervention coverage, morbidity, and mortality in Africa: a geospatial modelling analysis, Lancet Infect Dis 21(1) (2021) 59–69.

[32] G. Asmare, Willingness to accept malaria vaccine among caregivers of under-5 children in Southwest Ethiopia: a community based cross-sectional study, Malar J 21(1) (2022) 146.

[33] S.G. Smith, M.M. Zhou, The Bromodomain: A New Target in Emerging Epigenetic Medicine, ACS Chem Biol 11(3) (2016) 598–608.

[34] E.K.a.K. Henrick, Protein structure comparison service PDBeFold at European Bioinformatics Institute.

[35] G.A. Josling, S.A. Selvarajah, M. Petter, M.F. Duffy, The role of bromodomain proteins in regulating gene expression, Genes (Basel) 3(2) (2012) 320–43.

[36] P. Filippakopoulos, S. Knapp, The bromodomain interaction module, FEBS Lett 586(17) (2012) 2692–704.

[37] J. Moriniere, S. Rousseaux, U. Steuerwald, M. Soler-Lopez, S. Curtet, A.L. Vitte, J. Govin, J. Gaucher, K. Sadoul, D.J. Hart, J. Krijgsveld, S. Khochbin, C.W. Muller, C. Petosa, Cooperative binding of two acetylation marks on a histone tail by a single bromodomain, Nature 461(7264) (2009) 664–8.

[38] B. Padmanabhan, S. Mathur, R. Manjula, S. Tripathi, Bromodomain and extra-terminal (BET) family proteins: New therapeutic targets in major diseases, J Biosci 41(2) (2016) 295–311.

[39] K.A. Matthews, K.M. Senagbe, C. Notzel, C.A. Gonzales, X. Tong, F. Rijo-Ferreira, N.V. Bhanu, C. Miguel-Blanco, M.J. Lafuente-Monasterio, B.A. Garcia, B.F.C. Kafsack, E.D. Martinez, Disruption of the Plasmodium falciparum Life Cycle through Transcriptional Reprogramming by Inhibitors of Jumonji Demethylases, ACS Infect Dis 6(5) (2020) 1058–1075.

[40] M.J. Chua, D. Robaa, T.S. Skinner-Adams, W. Sippl, K.T. Andrews, Activity of bromodomain protein inhibitors/binders against asexual-stage Plasmodium falciparum parasites, Int J Parasitol Drugs Drug Resist 8(2) (2018) 189–193.

[41] D. Oladejo, G. Oduselu, T. Dokunmu, I. Isewon, E. Okafor, E.E.J. Iweala, E.F. Adebiyi, In Silico Evaluation of Inhibitors of Plasmodium Falciparum AP2-I Transcription Factor and Plasmodium Falciparum Bromodomain Protein 1, Research Square (2022).

[42] T. Hollin, M. Gupta, T. Lenz, K.G. Le Roch, Dynamic Chromatin Structure and Epigenetics Control the Fate of Malaria Parasites, Trends Genet 37(1) (2021) 73–85.

[43] C.G. Toenhake, S.A. Fraschka, M.S. Vijayabaskar, D.R. Westhead, S.J. van Heeringen, R. Bartfai, Chromatin Accessibility-Based Characterization of the Gene Regulatory Network Underlying Plasmodium falciparum Blood-Stage Development, Cell Host Microbe 23(4) (2018) 557–569 e9.

[44] L. Cui, J. Miao, T. Furuya, X. Li, X.Z. Su, L. Cui, PfGCN5-mediated histone H3 acetylation plays a key role in gene expression in Plasmodium falciparum, Eukaryot Cell 6(7) (2007) 1219–27.

[45] H.D. etal, Plasmodium bromodomain PfBDP4; A Target EnablingPackage, (2018).

[46] L.J. Clos, 2nd, M.F. Jofre, J.J. Ellinger, W.M. Westler, J.L. Markley, NMRbot: Python scripts enable high-throughput data collection on current Bruker BioSpin NMR spectrometers, Metabolomics 9(3) (2013) 558–563.

[47] W. Kabsch, Integration, scaling, space-group assignment and post-refinement, Acta Crystallogr D Biol Crystallogr 66(Pt 2) (2010) 133–44.

[48] A.J. McCoy, R.W. Grosse-Kunstleve, P.D. Adams, M.D. Winn, L.C. Storoni, R.J. Read, Phaser crystallographic software, J Appl Crystallogr 40(Pt 4) (2007) 658–674.

[49] P.D. Adams, P.V. Afonine, G. Bunkoczi, V.B. Chen, I.W. Davis, N. Echols, J.J. Headd, L.W. Hung, G.J. Kapral, R.W. Grosse-Kunstleve, A.J. McCoy, N.W. Moriarty, R. Oeffner, R.J. Read, D.C. Richardson, J.S. Richardson, T.C. Terwilliger, P.H. Zwart, PHENIX: a comprehensive Python-based system for macromolecular structure solution, Acta Crystallogr D Biol Crystallogr 66(Pt 2) (2010) 213–21.

[50] W.L. DeLano, The PyMOL Molecular Graphics System in, DeLano Scientific, Palo Alto, CA (2002).

