## Supplemental for "Structural insights into acetylated histone ligand recognition by the BDP1 bromodomain of *Plasmodium falciparum*"

### Supporting Information

[1]Department of Pharmacology, Larner College of Medicine, University of Vermont, Burlington, VT, 05405, USA

[2]Department of Pharmaceutical Sciences, Albany College of Pharmacy and Health Sciences, Colchester, VT, 05446, USA

[3]NMRFAM and Department of Biochemistry, University of Wisconsin-Madison, Madison, Wisconsin, 53706, USA

[4]Molecular Biology Consortium, Advanced Light Source, Berkeley, CA, 94720, USA

*Corresponding author

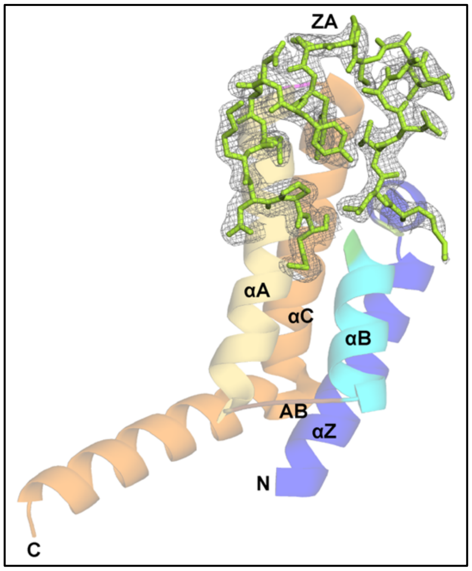

**Suppl. Fig. 1. Crystal structure of PfBDP1-BRD at 2.0 Å.** The $2F_O$ - $F_C$ electron density map for ZA loop in PfBDP1-BRD is displayed at 1.0σ contour level.

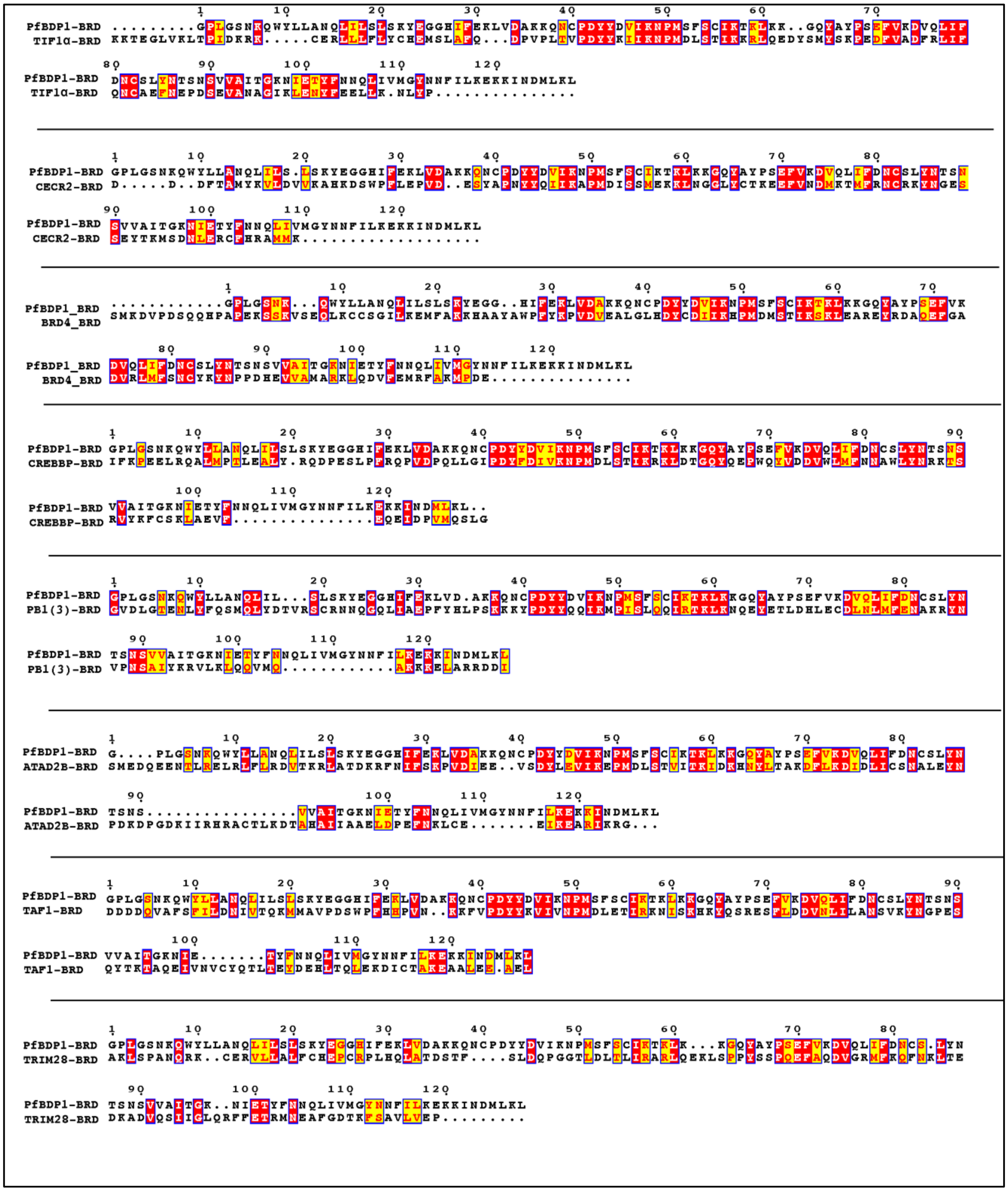

**Suppl. Fig. 2A. Sequence alignment of PfBDP1-BRD with human bromodomain.** The sequence alignment of PfBDP1-BRD with human bromodomain TIF1α (PDB ID: 3O33), CECR2 (PDB ID: 3NXB), BRD4 (PDB ID: 2OUO), CREBBP (PDB ID: 3DWY), PB1(3) (PDB ID: 3K2J), ATAD2B (PDB ID: 3LXJ), TAF1 (PDB ID: 3UV5) and TAF1 (PDB ID: 2RO1) was done using CLUSTAL W (1.83) multiple sequence alignment [1]. Sequence alignment figures were made using ESPript 3.0 [2]

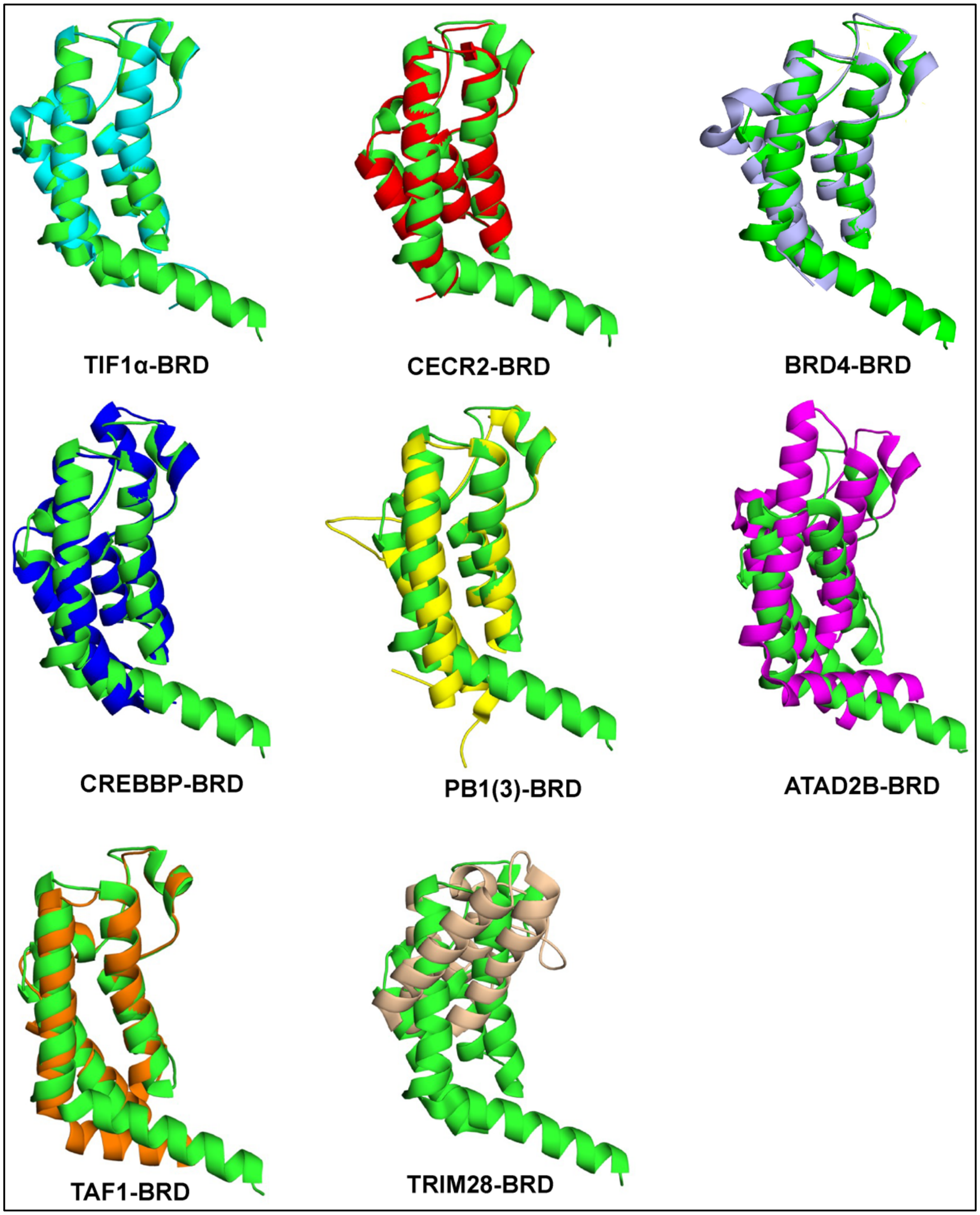

**Suppl. Fig. 2B. Structural alignment of PfBDP1-BRD with human bromodomain.** The structural alignment of PfBDP1-BRD (green) with human bromodomain TIF1α (PDB ID: 3O33 cyan), CECR2 (PDB ID: 3NXB red), BRD4 (PDB ID: 2OUO), CREBBP (PDB ID: 3DWY blue), PB1(3) (PDB ID: 3K2J yellow), ATAD2B (PDB ID: 3LXJ magenta), TAF1 (PDB ID: 3UV5 orange) and TAF1 (PDB ID: 2RO1 brown) was done using PyMOL [3] and the RMSD was calculated using the PDBeFold online server [4]

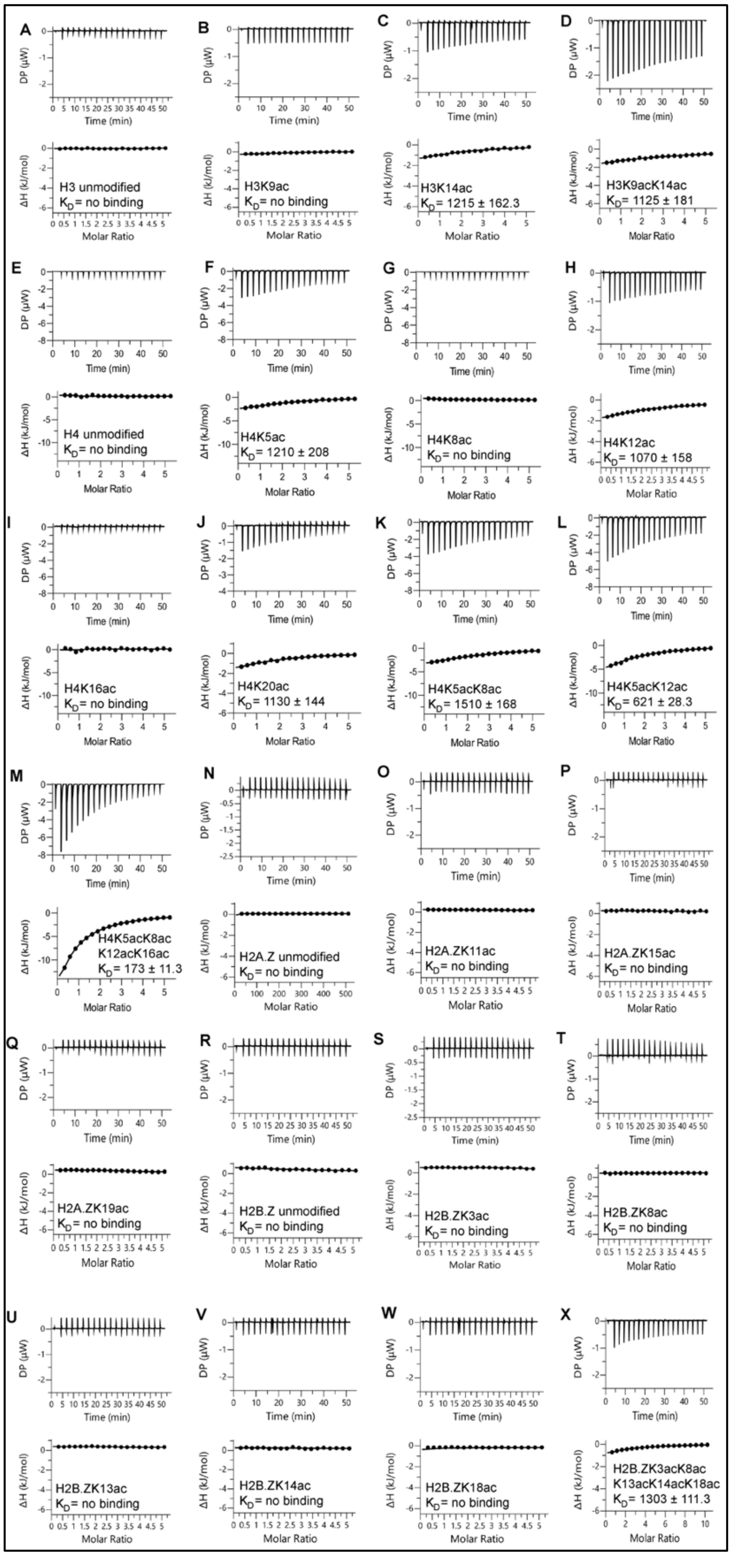

**Suppl. Fig. 3. ITC enthalpy plots for the binding of PfBDP1-BRD with acetylated histone peptides.** (A-D) PfBDP1-BRD with acetylated histone H3 peptides. (E-M) PfBDP1-BRD with acetylated histone H4 peptides. (N-Q) PfBDP1-BRD with acetylated histone H2A.Z peptides. (R-X) PfBDP1-BRD with acetylated histone H2B.Z peptides

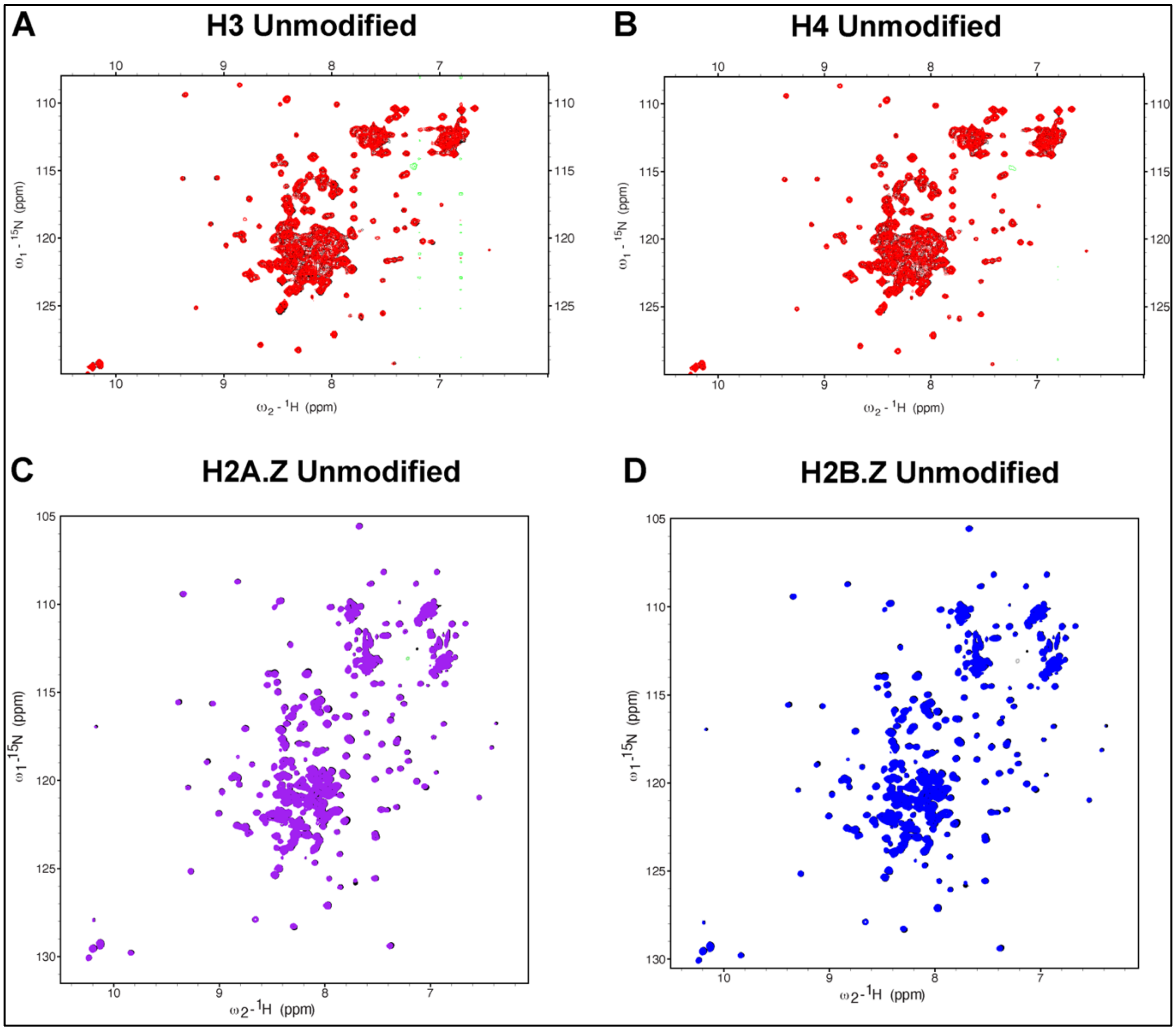

**Suppl. Fig. 4. Interaction of PfBDP1-BRD with unmodified histone peptide.** Superimposed 2D $^{15}N$-$^{1}H$ HSQC spectra of the $^{15}N$-labeled PfBDP1-BRD collected in titration experiments with the indicated histone peptides. The peaks of the PfBDP1-BRD apo protein are shown in black. The titration of PfBDP1-BRD with histone peptides was performed at 1:0 molar ratio (black), and in 1:5 molar ratio (red A-B, purple C and blue D). Each spectrum is labeled with the histone peptide used for the titration.

**References**

[1] R. Chenna, H. Sugawara, T. Koike, R. Lopez, T.J. Gibson, D.G. Higgins, J.D. Thompson, Multiple sequence alignment with the Clustal series of programs, Nucleic Acids Res 31(13) (2003) 3497-500.
[2] X. Robert, P. Gouet, Deciphering key features in protein structures with the new ENDscript server, Nucleic Acids Res 42(Web Server issue) (2014) W320-4.
[3] W.L. DeLano, The PyMOL Molecular Graphics System in, DeLano Scientific, Palo Alto, CA (2002).
[4] E.K.a.K. Henrick, Protein structure comparison service PDBeFold at European Bioinformatics Institute.
